# Ultrasensitivity and bistability in covalent modification cycles with positive autoregulation

**DOI:** 10.1101/2020.12.24.424291

**Authors:** C. Jeynes-Smith, R. P. Araujo

**Affiliations:** School of Mathematical Sciences, Queensland University of Technology, Brisbane, Australia; Institute of Health and Biomedical Innovation (IHBI), Brisbane, Australia

**Keywords:** chemical reaction networks, enzymes, posttranslational modifications, reaction kinetics, positive autoregulation

## Abstract

Switch-like behaviours in biochemical networks are of fundamental significance in biological signal processing, and exist as two distinct types: ultra-sensitivity and bistability. Here we propose two new models of a reversible covalent-modification cycle with positive autoregulation (PAR) - a motif structure that is thought to be capable of both ultrasensitivity and bistability in different parameter regimes. These new models appeal to a modelling framework that we call *complex complete*, which accounts fully for the molecular complexities of the underlying signalling mechanisms. Each of the two new models encodes a specific molecular mechanism for PAR. We demonstrate that the modelling simplifications for PAR models that have been used in previous work, which rely on a Michaelian approximation for the enzyme-mediated reactions, are unable to accurately recapitulate the qualitative signalling responses supported by our ‘full’ complex-complete models. Strikingly, we show that the parameter regimes in which ultrasensitivity and bistability obtain in the complex-complete framework contradict the predictions made by the Michaelian simplification. Our results highlight the critical importance of accurately representing the molecular details of biochemical signalling mechanisms, and strongly suggest that the Michaelian approximation may be inadequate for predictive models of enzyme-mediated chemical reactions with added regulations.

## 1. Introduction

The capacity for collections of biochemical reactions to respond in a switch-like, or *all-or-none*, manner is of fundamental significance in biological signal processing, and has been widely observed in many different signalling contexts [1–3]. Biochemical switch-like responses can exist as two distinct types [4,5]: ultrasensitivity, which is a steeply sigmoidal, monostable, dose-response profile; and bistability, a commonly-observed form of multistationarity, which constrains the system to exist in one of two possible stable steady-states for a given level of stimulus.

Ultrasensitivity has been implicated in functionally important switching mechanisms across a wide variety of biological contexts, ranging from cell signalling pathways [6,7], budding in yeast [8,9], maturation of xenopus oocytes [10,11], and embryonic differentiation [12]. More recently, ultrasensitivity has been recognised as an essential component in many robust perfect adaptation (RPA) mechanisms [13–15]. RPA is the ability of a system to asymptotically track a fixed ‘set-point’ following persistent perturbations to its interacting elements. This keystone signalling response is thought to be an essential characteristic of all evolvable and self-regulating biosystems [14,15], and has been ubiquitiously observed across all levels of biological organisation, from chemotaxis in single-celled organisms [16–27], to complex sensory systems [28–38]. Importantly, the dysregulation of RPA is believed to be linked to disorders such as cancer progression, drug addiction, chronic pain, and metabolic syndrome [39–44].

While the history of mathematical approaches to the study of ultrasensitivity-generating mechanisms may be traced back to the work of AV Hill [45] on cooperative binding in haemoglobin, the seminal work of Goldbeter and Koshland [46] on *zero-order* ultrasensitivity has exerted a profound influence on our modern understanding of ultrasensitivity. Indeed, it was Goldbeter and Koshland [46] who formally defined an ultrasensitive response as ‘an output response that is more sensitive to change in stimulus than the hyperbolic (Michaelis-Menten) equation’. Their landmark findings demonstrated that enzyme-mediated covalent modification cycles (such as phosphorylation/dephosphorylation cycles, or methylation/demethylation cycles, for instance) are capable of generating sensitivities comparable to cooperative enzymes with high Hill coefficients when the interconverting enzymes operate in their ‘zero-order’ regions (ie. in the region of saturation with respect to their protein substrates). Importantly, this was the first known example of *unlimited* ultrasensitivity: unlike the response of cooperative enzymes, where the steepness of the dose-response profile was limited, the zero-order mechanism allowed steepness to be increased indefinitely by ‘tuning’ certain parameter groups. More recently, a number of additional ultrasensitivity-generating mechanisms have been identified [47, 48], including inhibitor-generated ultrasensivity and substrate competition [48–51], cooperative binding [52,53], positive feedback [3,6,54], multisite phosphorylation [9,12,55], and negative feedback [8,48]. Some mathematical models also support the idea that cascade structures can promote ultrasensitivity [11].

In contrast to ultrasensitivity, bistability gives rise to a discontinuous switching mechanism. Bistability is widely believed to play a vital functional role in gene regulatory networks [56–60], cell differentiation [61,62], cell-cycle regulation [63,64], lineage commitment during development [2], exit from quiescence in mammalian cells [65], and biochemical and working memory [66,67]. Moreover, a bistable switch can exist as a one-way (irreversible) switch, or a two-way (‘toggle’) switch. Toggle switches are reversible but exhibit hysteresis, whereby once the switch has been ‘tripped’ by a suitable increase in stimulus, propelling the system to its upper steady-state, a *much larger decrease* in input stimulus is required to bring the system back down to its lower steady-state - a functionally important response in the context of fluctuating inputs [2,66,68]. One-way switches, on the other hand, can never return the system to its lower steady-state once the switch has been tripped. The aberrant formation of such one-way switches is thought to play a particularly pernicious role in cancer signalling dysregulation, by driving the constitutive activation of oncoproteins [5,69,70]. Mathematically, bistability is thought to arise from a variety of underlying mechanisms, including: positive autoregulation (PAR) and positive feedback, where a molecule either directly or indirectly promotes its own creation [56,57,59,65,66,71–73]; cooperative binding, where the binding of one molecule enhances the binding of subsequent molecules [63,74], antagonism, where one molecule benefits at the loss to another [62,75], and double negative feedback, a cycle in which two interacting molecules mutually inhibit each other [73].

Although emergent network behaviours such as bistability and ultrasensitivity, and the related phenomenon of RPA, have been the focus of many mathematical models, the detailed molecular mechanisms that might support these functions in ‘real’, highly complex signalling networks are yet to be understood. Indeed, a frontier research problem in biochemical signalling is the question of how specialised signal processing functions such as ultrasensitivity are implemented at the *microscale* of complex networks, through the highly intricate interactions of collections of proteins [15]. Even small collections of relatively simple enzyme-mediated reactions can involve the formation of a number of transient, intermediate molecular states, and can quickly give rise to complicated mathematical descriptions when all such states are explicitly accounted for. As a consequence, mathematical models of biochemical signal processing to date have typically been highly simplified - either by neglecting many of the proteins known to be involved, and frequently by simplifying the mathematical nature of their interactions - most notably through the (often indiscriminate) use of the Michaelis-Menten equation, involving the quasi-steady-state assumption (QSSA) on intermediate protein-protein complexes. Well-established and influential models of RPA in particular, have relied on such Michaelian descriptions of protein-protein interactions to suggest that specific parameter regimes promote robust adaptation by conferring ultrasensitivity at certain embedded ‘nodes’ of the network. The three-node RPA models published by Ma et al. [13], in particular, suggest that Michaelian models of positive autoregulation (PAR) are supportive of ultrasensitivity, and hence RPA, in very specific parameter regimes. But to what extent might these responses still be realized in more complex but realistic signalling descriptions, which account for all intermolecular interactions and intermediate protein-protein complexes?

In the present paper, we attempt to offer important new insights into this vitally important question. We propose several new, simple, yet detailed, mathematical models of covalent modification cycles incorporating PAR, appealing to a mathematical framework we call *complex-complete*. In this *complex-complete* framework, the molecular details of each specific PAR mechanism will be considered explicitly, and all intermediate protein-protein complexes accounted for. We will consider the capacity of each such model to exhibit bistability and ultrasensitivity, through model simulations over a wide range of parameters, and compare these outcomes to the behaviour supported by the well-established Michaelian model of PAR [13].

As a backdrop to these novel complex-complete models of PAR, we first briefly review the two main classes of simplified frameworks for the mathematical modelling of covalent modification cycles: simplified mass-action models, and Michaelian models (Fig 1).

**Figure 1.**
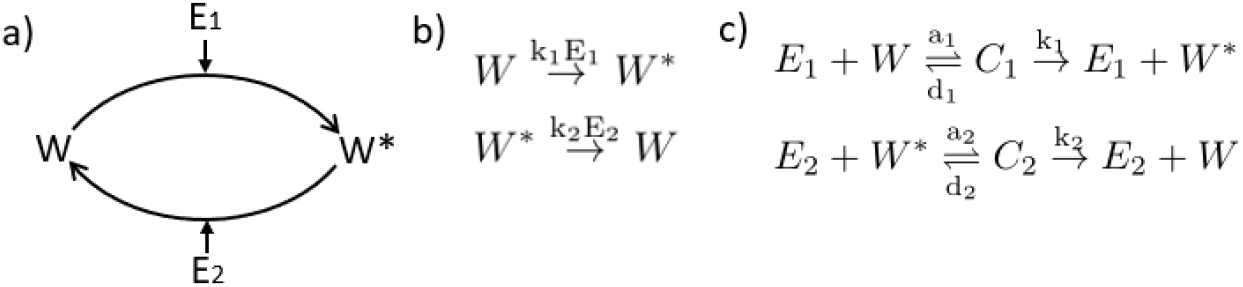
A mathematical framework for the modelling of a single covalent-modification cycle (a) A schematic representation of a reversible covalent-modification cycle in which a protein substrate (*W*) is chemically modified by an enzyme, *E*_1_, into an active form (*W*\*). The active form is, in turn, converted back to its unmodified form through the action of a second, independent, enzyme, *E*_2_. These processes may then be realised as a chemical reaction mechanism (b and c), with a corresponding graph structure, from which rate equations (ODEs) are ‘induced’ [76]. (b) A ‘complex-free’ mechanism, in which enzyme concentrations are incorporated into the biochemical rate constants (*k*_1_, *k*_2_) can be used as the basis of a simplified mass-action model of the cycle. (c) A reaction mechanism that explicitly includes transient enzyme-substrate complexes (*C*_1_, *C*_2_) can be used as the basis of either a *complex-complete* mass action model, or a Michaelian model. The Michaelian model is a simplification of the complex-complete framework, whereby complexes are assumed to reach a steady-state on a much faster time-scale than the other processes in the reaction, and whose steady-state concentrations are assumed to be negligible.

### (a) Simplified models of covalent modification cycles

Simplified mass-action models exclude intermediate protein-protein complexes (see Fig 1b). A simplified mass-action model of a covalent-modification cycle thus produces a single ordinary differential equation (ODE) for the output *W*\* (*t*), as in Eq (1.1),

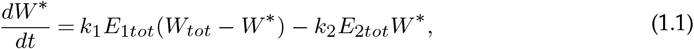

with

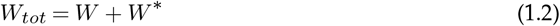

as a *conservation relation*, yielding the concentration of the unmodified protein, *W* (*t*), once *W*\* (*t*) is known from (1.1).

By contrast, Michaelis-Menten kinetics do take into account the existence of intermediate complexes in the system (see Fig 1c), but their explicit consideration is avoided through the application of the quasi steady-state assumption (QSSA). By the QSSA, complex association (*a*_*i*_) and dissociation (*d*_*i*_) processes are assumed to occur on a significantly faster timescale than the catalysis (*k*_*i*_) reaction. Moreover, the steady-state concentrations of the complexes (there being two enzyme-substrate complexes, *C*_1_ and *C*_2_, in a single covalent-modification cycle), having rapidly been reached, are assumed to be negligible in comparison with the total protein concentration (ie. *W*_*tot*_ = *W*\* + *W* + *C*_1_ + *C*_2_ ≈ *W*\* + *W*). We thereby obtain a single equation for *W*\*(*t*) (Eq (1.3)), which features parameter groups known as Michaelis constants, 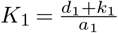 and 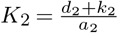,

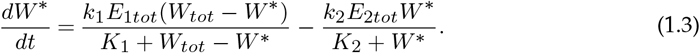

The first of the following conservation relations

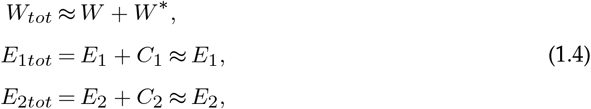

determines the concentration of *W* (*t*) once *W*\*(*t*) is established from (1.3).

We refer to models that consider all signalling events and intermediate molecules explicitly, thus providing a detailed representation of the signalling mechanisms, as *complex-complete mass-action models*. The complex-complete mass-action model associated with the chemical reaction network in Fig 1c can be expressed as a system of four ODEs and three conservation equations,

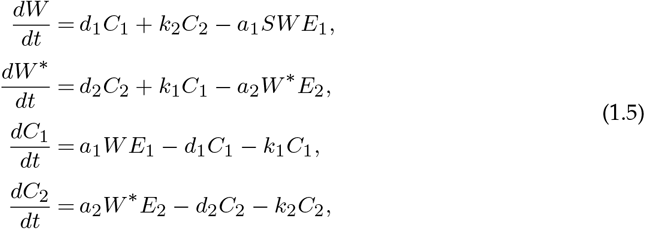

and

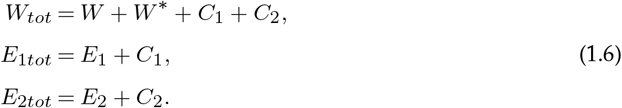

As mathematical models grow in size and complexity, simplified mass-action and Michaelian descriptions of the various components of the system can significantly reduce the number of variables and parameters to be considered. On the other hand, these benefits come at the expense of neglecting signalling events which could potentially alter the qualitative nature of the systems response in significant ways ([15,75,77]). Importantly, Goldbeter and Koshland’s seminal work on zero-order ultrasensitivity [46] has suggested that the Michaelian model of a covalent modification cycle, Eq (1.3), provides a good approximation to the ultrasensitive behaviour of the complex-complete mass-action model *provided* that *E*_1*tot*_, *E*_2*tot*_ ≪ *W*_*tot*_ - a parametric condition unlikely to obtain in many signal transduction networks, or other highly complex bionetworks in nature [14]. Moreover, if additional regulations such as autoregulatory interactions are present in the system, with an inevitable increase in the number of intermediate protein-protein complexes, it is unclear if simplified Michaelian descriptions of the system would be capable of recapitulating the qualitative behaviours of the corresponding complex-complete models for *any* parameter regime.

Although the larger number of variables and parameters often necessitates numerical, rather than analytical, approaches, these models allow the complexity of the system to speak for itself, and can be used to verify the extent to which simplified models of signalling motifs can recapitulate the behaviours of more detailed, accurate and mechanistic mathematical descriptions.

We now proceed to compare three different models of a covalent-modification cycle with positive autoregulation (PAR) - the well-established Michaelian model studied by Ma et al. [13], along with two novel complex-complete mass-action models.

### (b) A Michaelian Model of Positive Autoregulation (PAR)

In order to generate ultrasensitivity (and hence RPA when suitably embedded into an appropriate network topology) via positive autoregulation (PAR) of a covalent modification cycle, Ma et al. [13] used an equation of the following form:

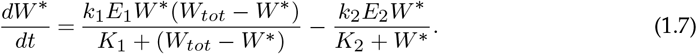

Here PAR has been incorporated into a Michaelian model by assuming that the rate of increase of *W*\* occurs in direct proportion to *E*_1_*W*\* (rather than just *E*_1_, as is the case for the Michaelian model without added regulations), thereby enabling the output molecule to enhance its own production. It is clear from the form of Eq (1.7) that if *K*_1_ ≪ *W*_*tot*_ − *W*\* and *K*_2_ ≫ *W*\*, then at steady state, 0 ≈ *W*\*(*k*_1_*E*_1_ − *k*_2_*E*_2_/*K*_2_). Thus, the system can exhibit unlimited ultrasensitivity as this parametric limit is approached, with a near-vertical dose-response occurring at 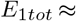 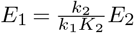.

In Fig 2 we examine how the Michaelis constants of this system are the key drivers of both bistability and (unlimited) ultrasensitivity in this model. In particular, for an initial choice of *K*_1_ ≈ *K*_2_ ≈ 1, the rate of increase in W* (orange/black lines), as given by the first term in Eq (1.7), intersects the rate of decrease (blue dashed line) in multiple places, indicating the existence of multiple steady states (bistability, in this case) for some values of *E*_1_. As we increase the value of *K*_2_ and decrease *K*_1_, we observe a narrowing in the range of values of *E*_1_ for which bistability obtains. This occurs as the negative rate of change approaches the same form as the positive rate of change. For *K*_1_ ≪ *W*_*tot*_ − *W*\* and *K*_2_ ≫ *W*\*, the two curves approach an exact match, and complete overlap, for 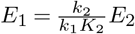. Under these conditions, every non-zero value of *W*\* is compatible with a steady state at 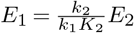 - a scenario that corresponds to unlimited ultrasensivity (ie. a vertical dose-response curve at the noted value of *E*_1_).

**Figure 2.**
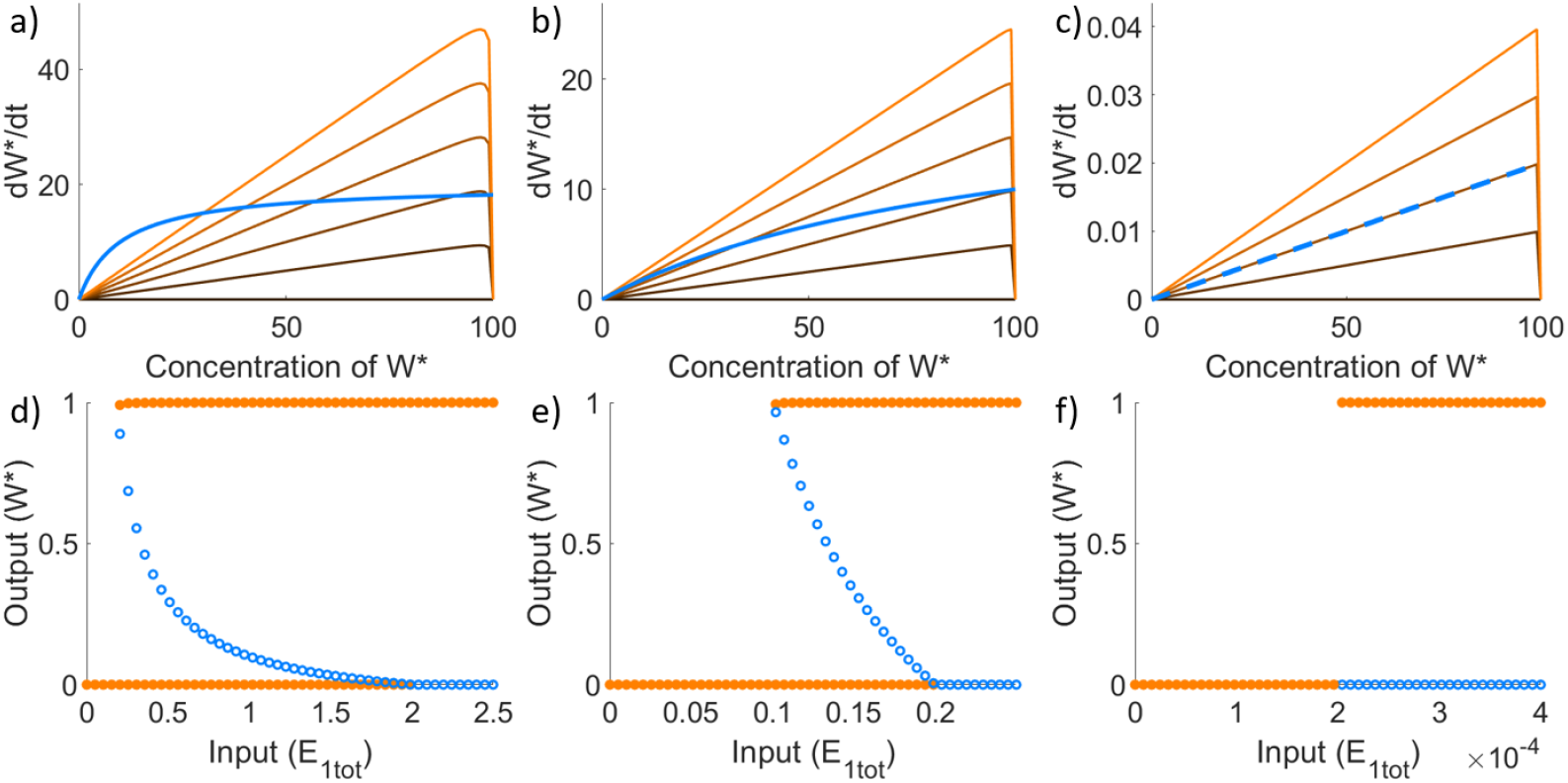
Bistability and Ultrasensitivity in a Michaelian model of PAR. The Michaelian model of PAR (Eq (1.7)) can exhibit both bistability, as a two-way (toggle) switch (a, b, d, e), and unlimited ultrasensitivity (c, f), depending on model parameters. Figures (a-c) plot the positive and negative contributions of the rate of change in *W*\* from Eq (1.7), for three different parameter sets. In each case, the negative component (independent of *E*_1_) is represented by the blue line. Since the positive component is a function of *E*_1_ (the input), we represent the positive component of the rate change by the solid line which changes from black to orange with an increasing value of *E*_1_. The intersection of the positive and negative components thus indicates the steady states of the system, which are plotted in (d-f), respectively, for a range values of *E*_1_. Stable steady states are represented by solid circles, with unstable states represented by open circles. As shown, figures (a) and (b) have multiple intersection points for certain values of *E*_1_. These existence of multiple steady states is highlighted by the bistable plots in the corresponding figures (d) and (e). Parameters: all six plots use *k*_1_ = *k*_2_ = 1, *W*_*tot*_ = 100, *E*_2_ = 20. In addition, (a) and (d) use: *K*_1_ = 10^−1^ and *K*_2_ = 10^1^; (b) and (e) use: *K*_1_ = 10^−2^ and *K*_2_ = 10^2^; and (c) and (e) use: *K*_1_ = 10^−5^ and *K*_2_ = 10^5^.

Note that in the Michaelian model of PAR by Ma et al. [13], there is no consideration of *how* the output molecule *W*\* might upregulate its own production through its interactions with other molecules, and thus no accounting for any particular autoregulatory *mechanism*. As a consequence, there is no way to determine how well this model might reflect PAR mechanisms in ‘real’ biochemical reaction networks. In order to consider the role of the *mechanism* in PAR, we propose two possible chemical reaction mechanisms - a direct positive autoregulatory mechanism (hereafter, PAR-d), and an indirect mechanism (hereafter, PAR-i).

### (c) A novel model of Direct Positive Autoregulation (PAR-d)

In order implement a direct PAR mechanism we supplement the basic covalent-modification cycle (Fig 1c) to include a third set of reactions whereby the output, *W*\*, binds in trans to the unmodified protein *W*, thereby catalysing the conversion of the latter to *W*\*. In the process of doing so, an intermediate protein-protein complex comprising both *W* and *W*\* is formed. We represent this process schematically in Fig 3a and by the chemical reaction network in Fig 3b.

**Figure 3.**
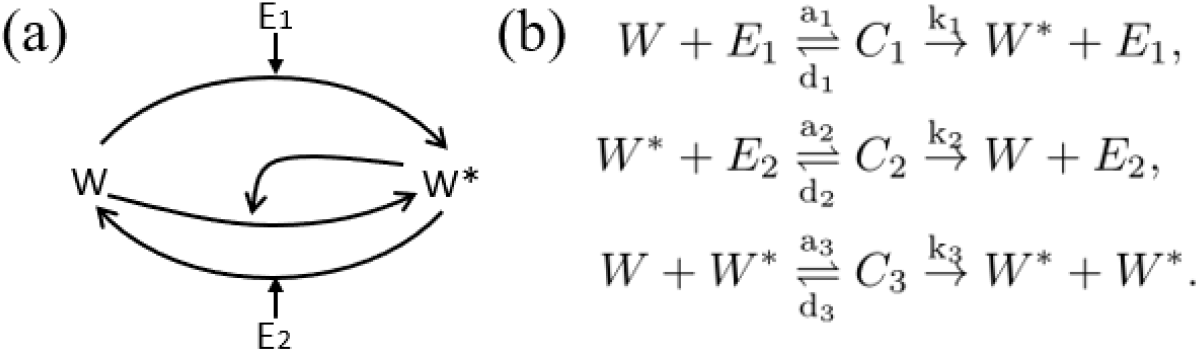
PAR-d System. (a) Schematic representation of the PAR-d mechanism. (b) Chemical reaction network corresponding to the PAR-d mechanism, comprising three groups of reactions: (i) *E*_1_ binding reversibly to *W*, forming a complex *C*_1_, and catalysing the production of *W*\* from *W*; (ii) *E*_2_ binding reversibly to *W*\*, forming a complex *C*_2_, and catalysing the production of *W* from *W*\*; and (iii) *W*\* binding reversibly to *W*, forming a complex *C*_3_, and catalysing the conversion of *W* to *W*\*.

This chemical reaction network, through the law of mass action, induces the following system of ordinary differential equations:

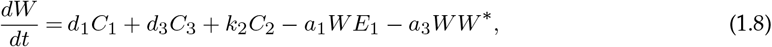

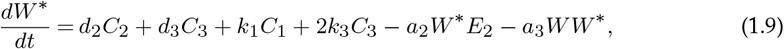

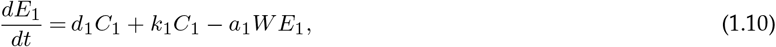

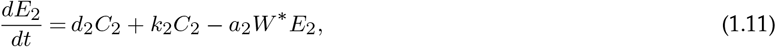

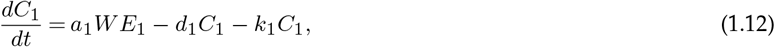

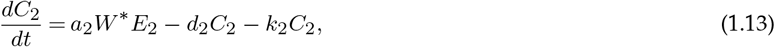

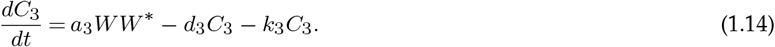

The summation of Eqs (1.8), (1.9), (1.12), (1.13), and (1.14) yields a conservation equation for the substrate protein,

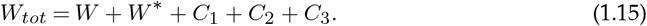

Likewise, summation of Eqs (1.10), and (1.12), and of Eqs (1.11), and (1.13), yield conservation equations for the two enzymes:

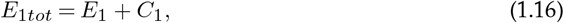

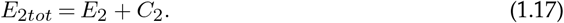

### (d) A novel model of Indirect Positive Autoregulation (PAR-i)

For the indirect implementation of PAR we introduce two new reaction groups to the reversible covalent-modification cycle. First, we add a reaction in which the output, *W*\*, can bind reversibly with the enzyme, *E*_1_, forming the transient complex *C*_3_. Then, in a second reaction group, the complex *C*_3_ can bind reversibly to *W*, forming another complex *C*_4_, which catalyses the conversion of *W* to *W*\*. In this way, the binding of *W*\* to *E*_1_ (producing *C*_3_) alters the efficiency of the enzyme *E*_1_ through an allosteric interaction. Thus, in contrast to PAR-d, where *W*\* catalyses its own production from the unmodified form *W*, *W*\* now enhances its own production through an indirect mechanism, through its interactions with *E*_1_. We illustrate the PAR-i mechanism by the schematic in Fig 4a, and produce a corresponding chemical reaction network in Fig 4b.

**Figure 4.**
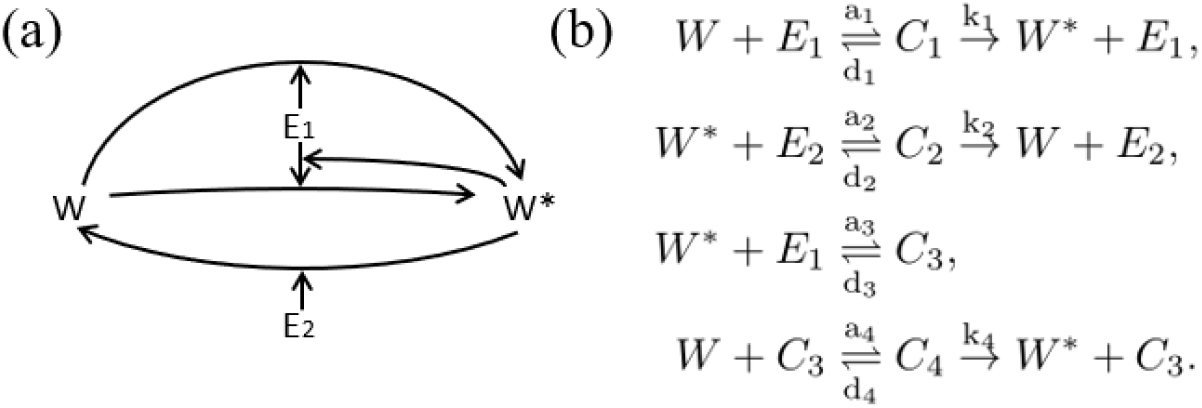
PAR-i System. (a) Schematic representation of the PAR-i mechanism. b) A chemical reaction network for PAR-i. This includes four reaction groups: (i) *E*_1_ binding reversibly to *W*, forming a complex *C*_1_, and catalysing the production of *W*\* from *W*; (ii) *E*_2_ binding reversibly to *W*\* forming a complex *C*_2_, and catalysing the production of *W* from *W*\*; (iii) *W*\* binding reversibly with *E*_1_ to form a complex *C*_3_; and (iv) *C*_3_ (containing enzyme *E*_1_) binding reversibly with *W*, forming a complex *C*_4_, catalysing the conversion of *W* to *W*\*.

By the Law of Mass-action, we obtain a system of ordinary differential equations from the chemical reaction network in Fig 4b:

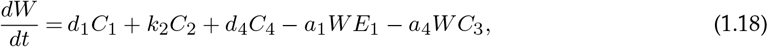

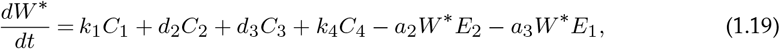

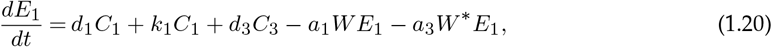

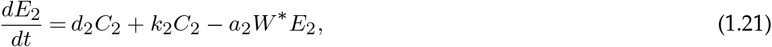

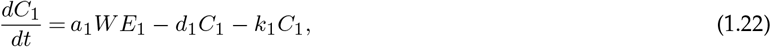

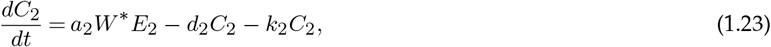

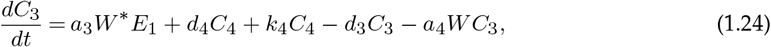

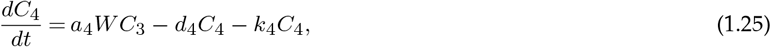

The summation of Eqs (1.18), (1.19), (1.22), (1.23), (1.24), and (1.25) gives a conservation equation for the substrate protein,

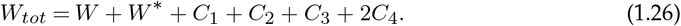

Similarly, the summation of Eqs (1.20), (1.22), (1.24), and (1.25), and of Eqs (1.21) and (1.23), yield conservation equations for the two interconverting enzymes:

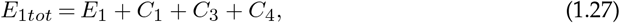

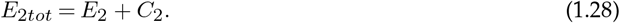

## 2. Results

We have undertaken extensive numerical explorations of the dose-response profiles of our two new complex-complete models of PAR, examining how the steady-state concentration of the output (*W*\*) changes with the input (*E*_1*tot*_). In all figures we show *W*\* as a proportion of the total substrate protein available in the system (*W*_*tot*_). These simulations consider the qualitative influence of each individual rate constant (*a*_*i*_, *d*_*i*_, *k*_*i*_), the total protein abundances (*W*_*tot*_ and *E*_2*tot*_, considered constants) as well as key parameter groups - the Michaelis constants (*K*_1_, *K*_2_, *K*_3_), and the dissociation constant (*K*_*d*_) that occurs in PAR-i. Each parameter, or parameter group, is varied in turn over the range (10^−10^, 10^10^) while holding all other parameters fixed. Linear stability analysis is used to classify the stability of all steady-states.

Our overarching goal is to determine the extent to which bistability (as a one-way switch and/or as a toggle switch) and ultrasensitivity are obtained in our complex-complete models of PAR, and to establish the similarities and differences in the predictions of these detailed models in comparison with the simpler Michaelian model of PAR.

### (a) PAR-d readily exhibits bistability and can create both one-way and two-way switches

Our extensive computational simulations reveal that the PAR-d model is highly prone to bistability. Indeed, whenever a monostable profile was obtained, almost any attempt to further increase the sensitivity (steepness of the slope) of the dose-response curve by increasing (or decreasing) a suitable parameter ultimately led to the onset of bistability.

Strikingly, the existence of bistability in the dose-response of PAR-d differs from that in the simple Michaelian model in a number of fundamental ways. First, unlike the Michaelian model which only admits two-way (toggle) switches in its bistable regime, PAR-d also has the potential for one-way (irreversible) switches. In Fig 5 we demonstrate how altering a single parameter can shift the system’s response from monostable ultrasensitivity, to a bistable two-way switch, to a bistable one-way switch.

**Figure 5.**
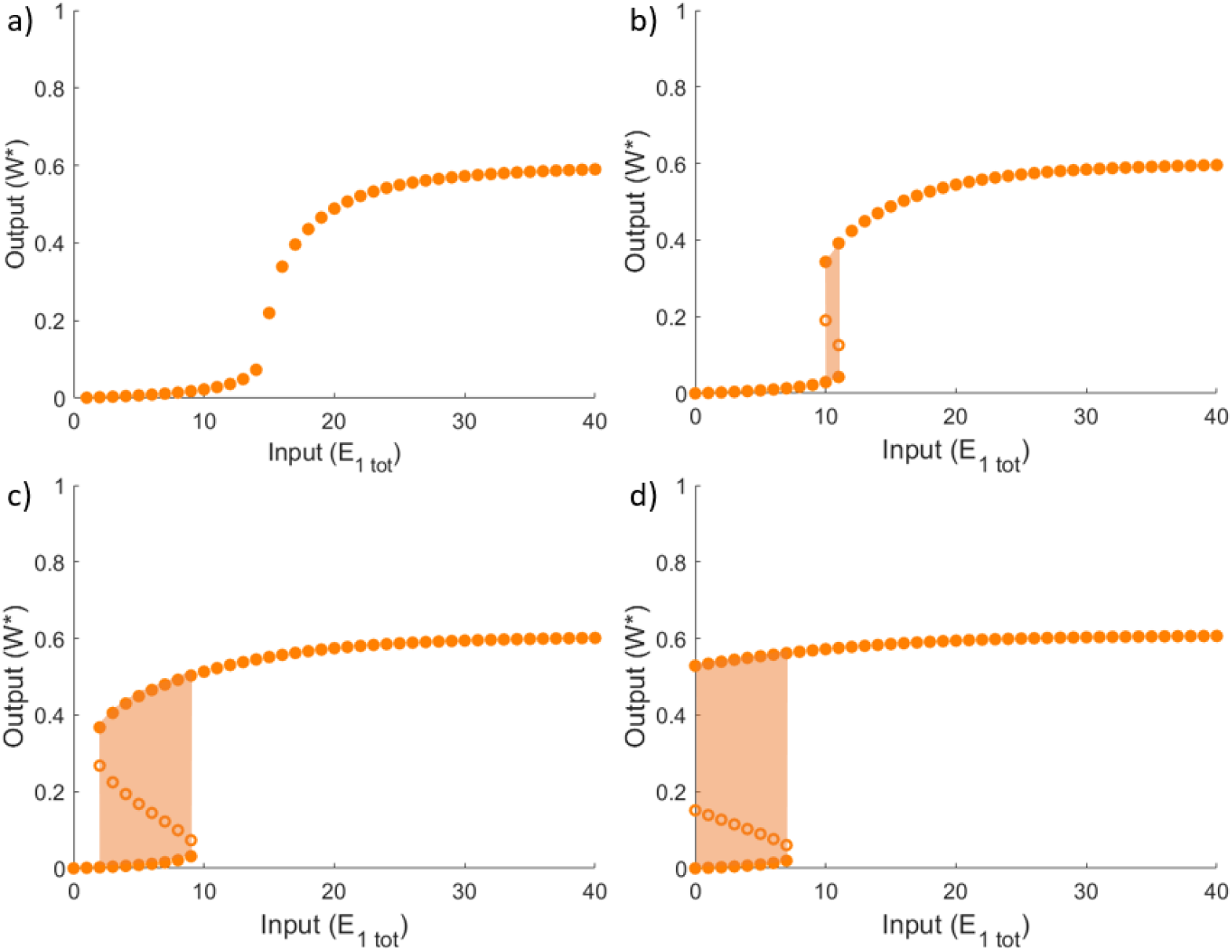
Bistability in PAR-d. Analysis of PAR-d dose-response profiles for (a) a continuous, ultrasensitive regime, a regime corresponding to a very narrow bistable regions, just past the bifurcation point that separates ultrasensitivity from bistability, (c) a bistable regime with a two-way switch, and (d) a bistable regime with a one-way switch. Solid circles denote stable solutions, while open circles denote unstable solutions. The shaded areas highlight the bistable regions. Parameters: All plots use *W*_*tot*_ = 100, *E*_2*tot*_ = 20, *a*_1_ = *d*_1_ = *k*_1_ = *a*_2_ = *d*_2_ = *k*_2_ = 1. The different profiles vary in the rate constants *a*_3_, *d*_3_, and *k*_3_: (a) *a*_3_ = 0.01, and *d*_3_ = *k*_3_ = 1; (b) *a*_3_ = 0.02, and *d*_3_ = *k*_3_ = 2; (c) *a*_3_ = 0.03, and *d*_3_ = *k*_3_ = 3 and d) *a*_3_ = 0.04, and *d*_3_ = *k*_3_ = 4.

Furthermore it is intriguing to observe that, in stark contrast to the Michaelian model, the capacity for bistability in PAR-d is *not* determined by the Michaelis constants alone. In Fig 5, for instance, a full range of behaviours is obtained from the model - from a monostable ultrasensitive profile, through to both one- and two-way switches - all generated with a single set of Michaelis constants *K*_1_, *K*_2_, *K*_3_. In fact, for each of the three individual rate constants, *a*_*i*_, *d*_*i*_, *k*_*i*_ that comprise a Michaelis constant, *K*_*i*_, we discovered that it is the catalytic constant, *k*_*i*_, specifically, that controls the qualitative response of the system and the shape of its dose-response profile.

We explore this finding further in Fig 6, where we examine the various possible combinations of holding a Michaelis constant fixed, along with *one* of its component rate constants, while varying the two remaining rate constants. In Fig 6a, we hold *K*_1_ and *k*_1_ constant, while increasing *d*_1_ and *a*_1_ in a proportion that maintains the constant *K*_1_. The profiles that are generated with these changes are all identical. By contrast, when we alter *k*_1_ and *a*_1_ (Fig 6b) and *k*_1_ and *d*_1_ (Fig 6c) we observe clear differences amongst the profiles. We also observe that across Fig 6b and Fig 6c, dose-response profiles associated with the matching values of *K*_1_ and *k*_1_ are the same irrespective of the value of *a*_1_ and *d*_1_. These same behaviours can be observed in the second (Fig 6d-f) and third (Fig 6g-i) reactions.

**Figure 6.**
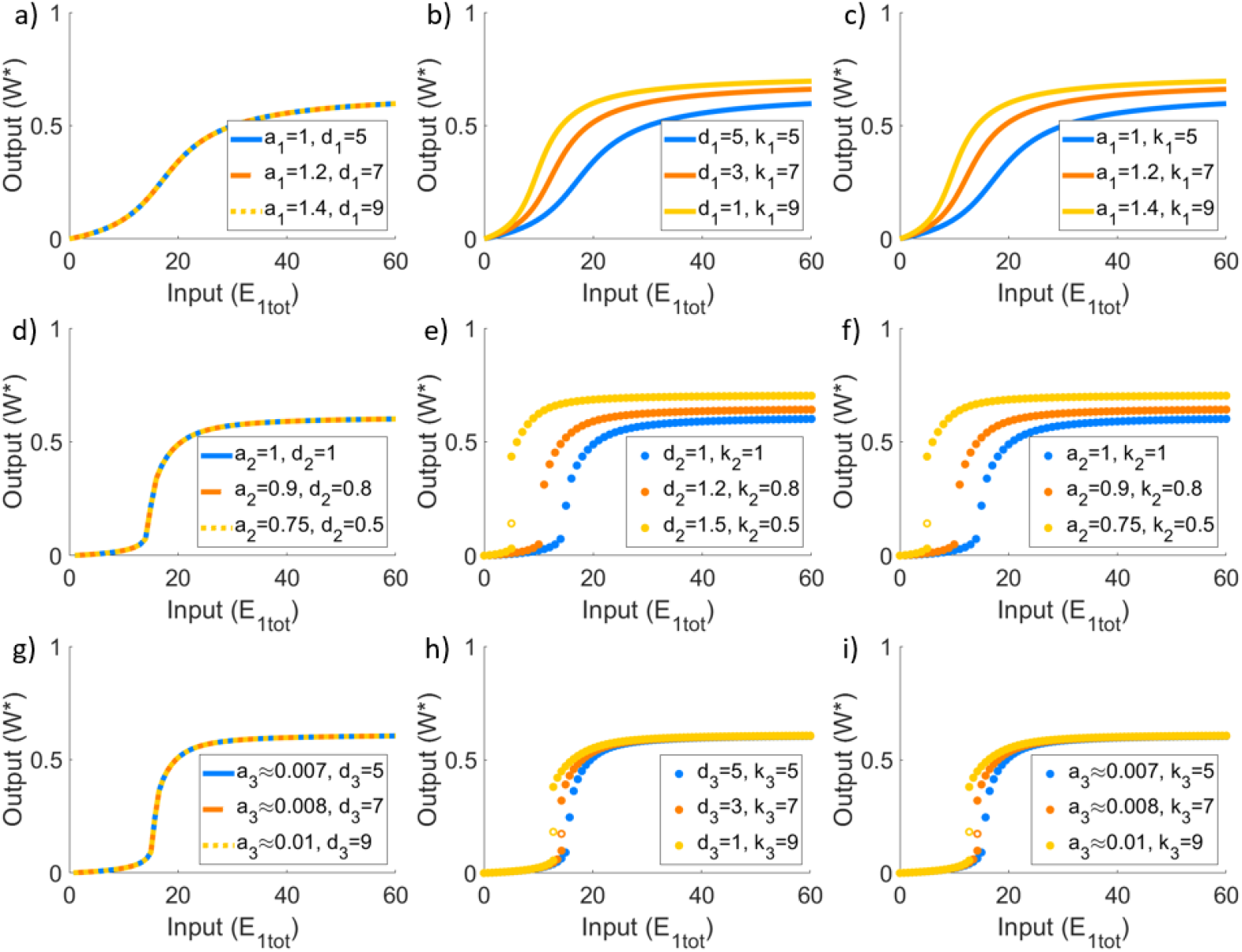
Varying rate constants in PAR-d. Pairs of independent rate constants are altered whilst holding the corresponding values of *K*_1_, *K*_2_, and *K*_3_ fixed. In (a), (d), (g) we find that holding the catalytic constant fixed results in no changes to the dose-response profile. Parameters: *W*_*tot*_ = 100 and *E*_2*tot*_ = 20 for all plots. For (a)-(c): *a*_1_ = *a*_2_ = *d*_3_ = *k*_3_ = 1, *d*_1_ = *k*_1_ = *d*_2_ = *k*_2_ = 5, *a*_3_ = 0.01; for (d)-(f): *a*_1_ = *d*_1_ = *k*_1_ = *a*_2_ = *d*_2_ = *k*_2_ = *d*_3_ = *k*_3_ = 1, *a*_3_ = 0.01; for (g)-(i): *a*_1_ = *d*_1_ = *k*_1_ = *a*_2_ = *d*_2_ = *k*_2_ = 1, *a*_3_ = 0.007, and *d*_3_ = *k*_3_ = 5. Varied rate constants identified in the figure legends. Closed circles indicate stable steady-states; open circles are unstable steady-states.

In fact, it is clear that the catalytic constants (*k*_*i*_) play a unique and important role in tuning the shape of the system’s dose-response profiles. In Figs 5 and 6, the parameters are altered to increase the catalytic constant in question without also altering the corresponding Michaelis constant. On the other hand, by increasing *k*_1_ on its own, and hence increasing *K*_1_ simultaneously, the system changes from monostable ultrasensitivity, to bistability, and then back to monostable ultrasensitivity (see Fig 7) - an unusual and unexpected behaviour not afforded by the simple Michaelian model. Remarkably, the monostable regions of these profiles continue to increase in sensitivity, as this complex phenomenon unfolds, despite the inclusion of the bistable region. This behaviour is tied uniquely and specifically to the first catalytic constant, *k*_1_.

**Figure 7.**
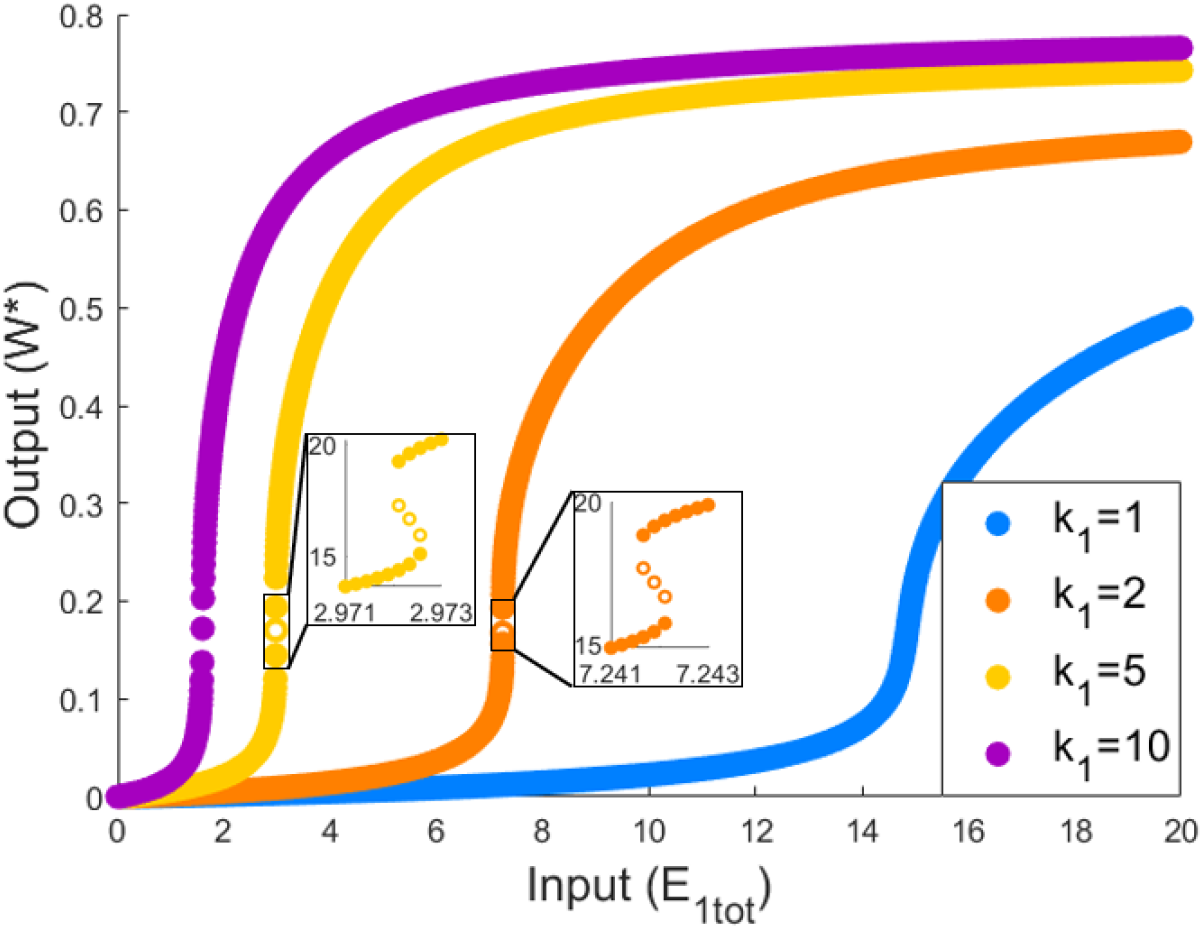
The role of the catalytic constant *k*_1_ in PAR-d. Increasing *k*_1_ while holding all other biochemical rate constants fixed engenders a complex response in the system’s dose-response profile. As illustrated here, continued increases in *k*_1_ transform the dose-response profile from monostable, to bistable, and back to monostable, all the while increasing the sensitivity of the monostable regions. Parameters: *a*_1_ = *d*_1_ = *a*_2_ = *d*_2_ = *k*_2_ = *d*_3_ = *k*_3_ = 1, and *a*_3_ = 0.01, *W*_*tot*_ = 100, *E*_2*tot*_ = 20. Closed circles indicate stable steady-states; open circles are unstable steady-states.

Moreover, while PAR-d can exhibit ultrasensitivity as demonstrated in Figs 6 and 7, the steepness of this near-vertical region of the dose-response cannot be increased indefinitely - in marked contradistinction to the Michaelian model which supports ulimited ultrasensitivity. As shown in Fig 7, increasing *k*_1_ can increase the sensitivity of the dose-response profile, but the steepness that can be achieved is limited. In addition, the location of the ultrasensitive portion of the curve is consistently moved to the left, towards the origin, which also constrains the development of ultrasensitivity due to this particular parameter change. In Fig 8, decreasing *K*_1_ with fixed *k*_1_ highlights the limit in ultrasensitivity that is approached as *K*_1_ is reduced.

**Figure 8.**
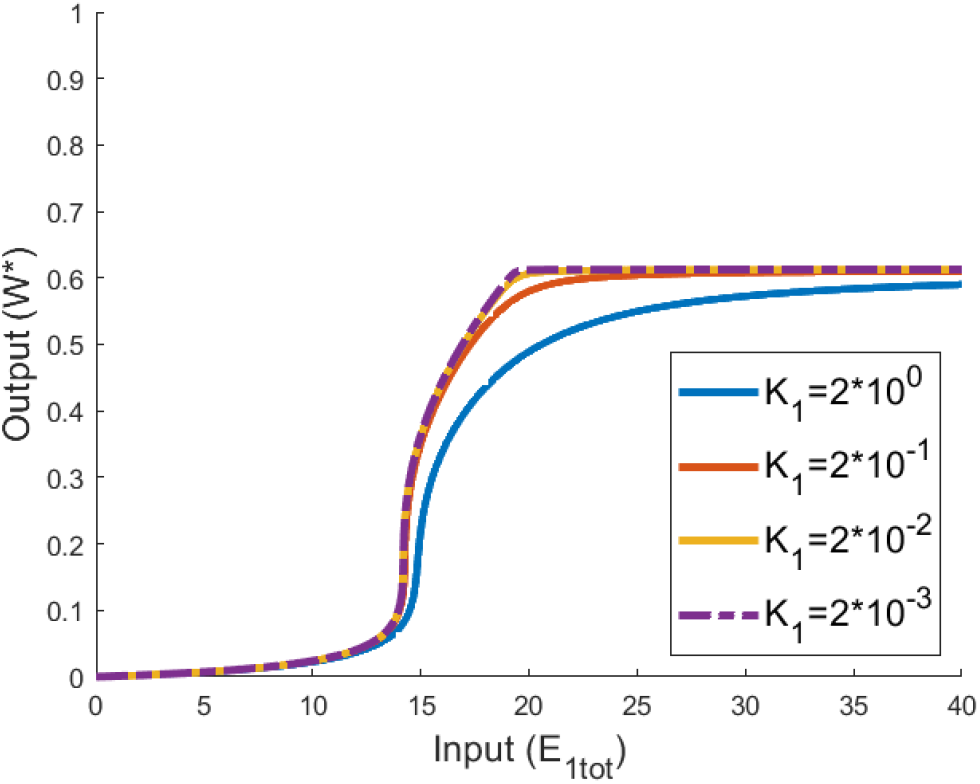
The role of Michaelis constant *K*_1_ in PAR-d. Here we decrease the value of *K*_1_ by increasing the value of *a*_1_ whilst holding *d*_1_ and *k*_1_ fixed. As shown, increasing *K*_1_ increases sensitivity without giving rise to bistability, but cannot increase sensitivity indefinitely. Sensitivity is clearly observed to approach a limiting state. Parameters: *W*_*tot*_ = 100, *E*_2*tot*_ = 20, *d*_1_ = *k*_1_ = *a*_2_ = *d*_2_ = *k*_2_ = *d*_3_ = *k*_3_ = 1, and *a*_3_ = 0.01. To achieve the indicated values of *K*_1_, we set *a*_3_ = 1 (blue), *a*_1_ = 10 (red), *a*_1_ = 100 (yellow), and *a*_1_ = 1000 (purple).

Altering the other parameters of the PAR-d model (*K*_2_, *K*_3_, and *W*_*tot*_), in an attempt to increase sensitivity, ultimately culminates in the onset of bistability. Unlike the situation illustrated in Fig 7, where the system switches from monostable to bistable and back to monostable for a monotone increase in a single parameter, bistability now persists once it is triggered at a critical value of the parameter in question. In particular, in Fig 9, we observe that decreasing *K*_2_ increases the sensitivity in the lower half of the substrate profile, without greatly affecting the sensitivity in the upper half. This ultimately leads to bistability as the lower half is shifted too far to the right. Further decreases in *K*_2_ do not lead to the creation of a one-way switch, as we illustrate by the profiles for *K*_2_ = 0.4 and *K*_2_ = 0.2 which can be observed to overlap without significant change.

**Figure 9.**
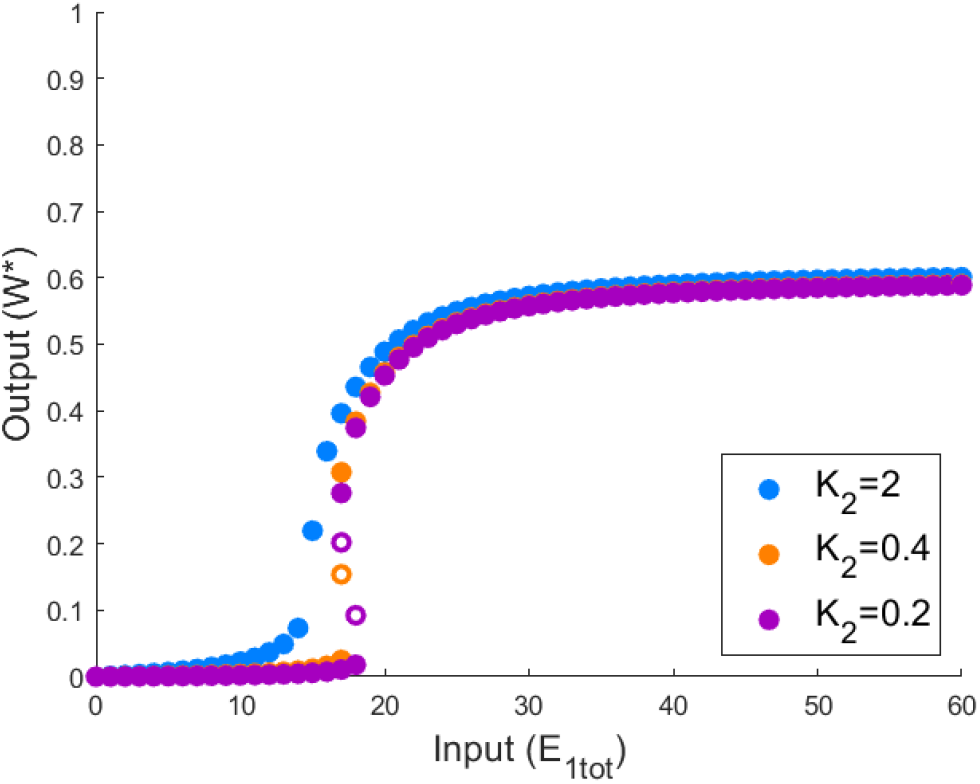
The role of Michaelis constant *K*_2_ in PAR-d. Here we alter the value of *K*_2_, by altering the value of *a*_2_ whilst holding *d*_2_ and *k*_2_ fixed. As shown, decreases in *K*_2_ ultlimately lead to bistability, but can only create a two-way (not one-way) switch. Parameters: *W*_*tot*_ = 100, *E*_2*tot*_ = 20, *a*_1_ = *d*_1_ = *k*_1_ = *d*_2_ = *k*_2_ = *d*_3_ = *k*_3_ = 1, and *a*_3_ = 0.01. To achieve the indicated values of *K*_2_, we set *a*_2_ = 1 (blue), *a*_2_ = 5 (orange), and *a*_2_ = 10 (purple). Solid circles indicate stable steady-states; open circles indicate unstable steady-states.

For the parameters *K*_3_ (Fig 10a) and *W*_*tot*_ (Fig 10b), on the other hand, continued monotone alterations in these parameters transform the two-way switch into a one-way switch. By either decreasing *K*_3_ or increasing *W*_*tot*_, the upper portion of the dose-response profile is shifted rapidly to the left, much more rapidly than the shifting of the lower portion, which leads to an overlap, creating bistability. This trend continues and ultimately leads to the creation of a one-way switch as the upper portion of the curve moves to the left past the location of the vertical axis.

**Figure 10.**
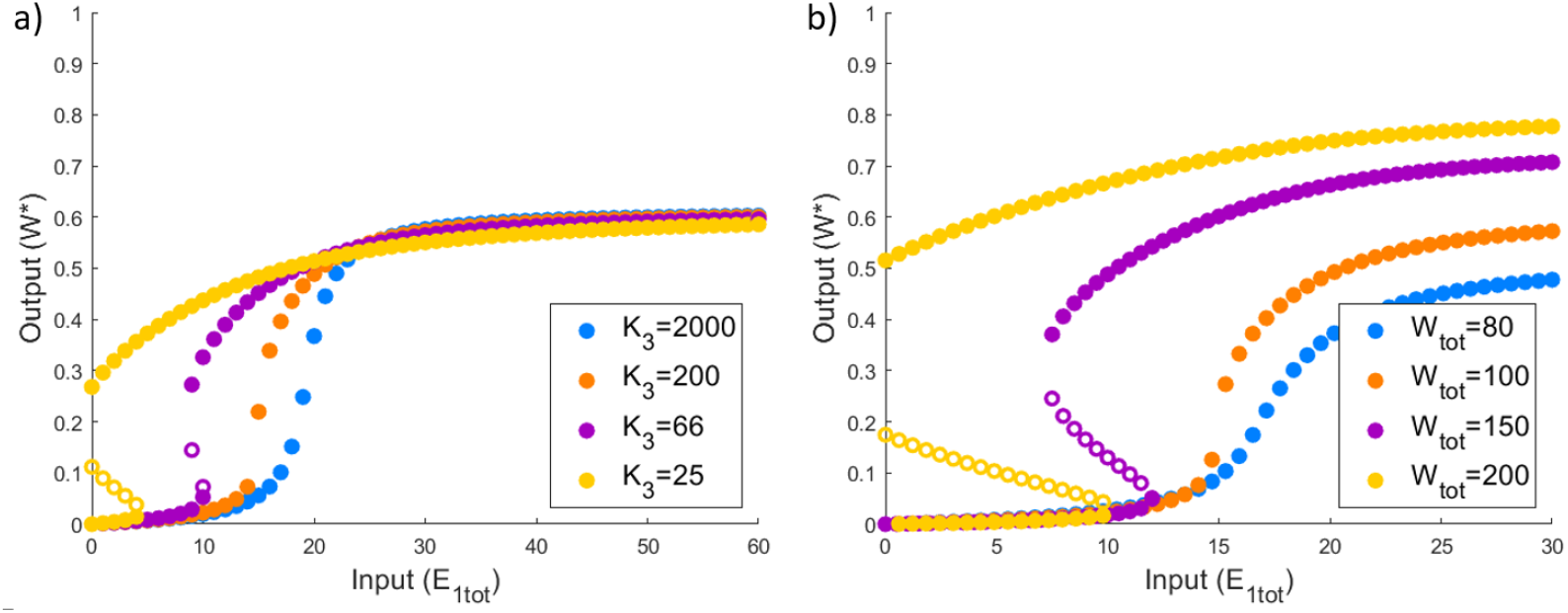
The role of Michaelis constant, *K*_3_, and protein abundance, *W*_*tot*_, in PAR-d. Continued decreases in *K*_3_, or increases in *W*_*tot*_ transform an initially monostable (ultrasensitive) profile into a bistable one, ultimately leading to the creation of a one-way switch. Parameters: *E*_2*tot*_ = 20, *a*_1_ = *d*_1_ = *k*_1_ = *a*_2_ = *d*_2_ = *k*_2_ = *d*_3_ = *k*_3_ = 1. (a) *W*_*tot*_ = 100, with *a*_3_ = 0.001 (blue), *a*_3_ = 0.01 (orange), *a*_3_ = 0.03 (purple), and *a*_3_ = 0.08 (yellow). (b) *a*_3_ = 0.01, with *W*_*tot*_ = 80 (blue), *W*_*tot*_ = 100 (orange), *W*_*tot*_ = 150 (purple), and *W*_*tot*_ = 200 (yellow).

### (b) PAR-i readily exhibits monostable responses but can produce two-way (*not* one-way) switches

Through extensive numerical simulations we find that PAR-i is tremendously less prone to bistability than the PAR-d model (which is highly prone to bistability) or the Michaelian model. Indeed, we find that the catalytic constant *k*_4_ is the *only* parameter in this model that drives bistability (see Fig 11). This is a key distinction from the Michaelian model, wherein the two Michaelis constants are the drivers of bistability, or the PAR-d model, where a number of parameters drive bistability as discussed in the previous section.

**Figure 11.**
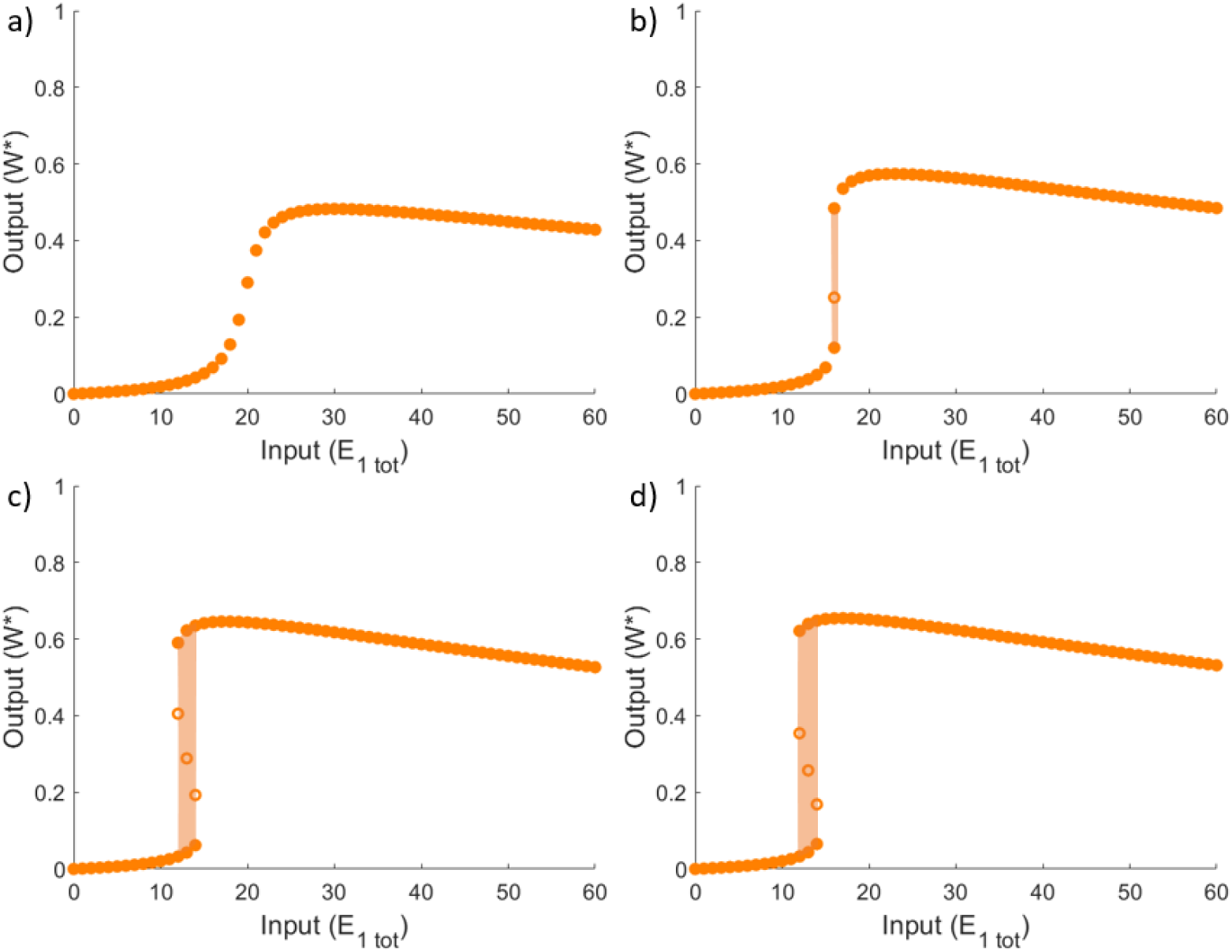
Bistability in PAR-i. Analysis of steady-state solutions of PAR-i model for (a) a continuous, monostable, ultrasensitive regime, (b) a bistable regime which is very close to the bifurcation between continuous ultrasensitivity and bistability, and (c)-(d) bistable regimes with a two-way switch. solid circles indicate stable solutions, empty circles indicate unstable solutions. Shaded areas highlight bistable the region. Parameters: all plots use *W*_*tot*_ = 100, *E*_2*tot*_ = 20, *a*_1_ = *a*_2_ = *a*_4_ = *d*_1_ = *k*_1_ = *d*_2_ = *k*_2_ = *d*_4_ = 1, *d*_3_ = 10, and *a*_3_ = 0.1. The different profiles are created by increasing the rate constant *k*_4_: (a) *k*_4_ = 1; (b) *k*_4_ = 3; (c) *k*_4_ = 30; and (d) *k*_4_ = 300.

The relationship between bistability and the parameter regime in the PAR-i model is quite subtle. In particular, while increasing the value of *k*_4_ is able to convert a sensitive but monostable profile into a bistable one, it is striking to observe that many other parameters can *predispose* the system to bistability, enabling the bifurcation to occur at a lower value of *k*_4_. In Fig 12, for instance, we show that for a given value of *k*_4_, we are able to switch between ultrasensitivity and bistability by increasing the sensitivity of the system via a reduction in the Michaelis constants. In particular, decreasing the values of *K*_2_ and *K*_4_ increases the sensitivity of the underlying ultrasensitive monostable profile, allowing bistability to be triggered for smaller values of *k*_4_. Interestingly, when *K*_1_ is decreased, the system is ‘pushed back’ towards monostability. Nevertheless, Michaelis constants are not considered drivers of bistability in the sense that, for sufficiently low values of *k*_4_, they are unable to achieve bistability on their own.

**Figure 12.**
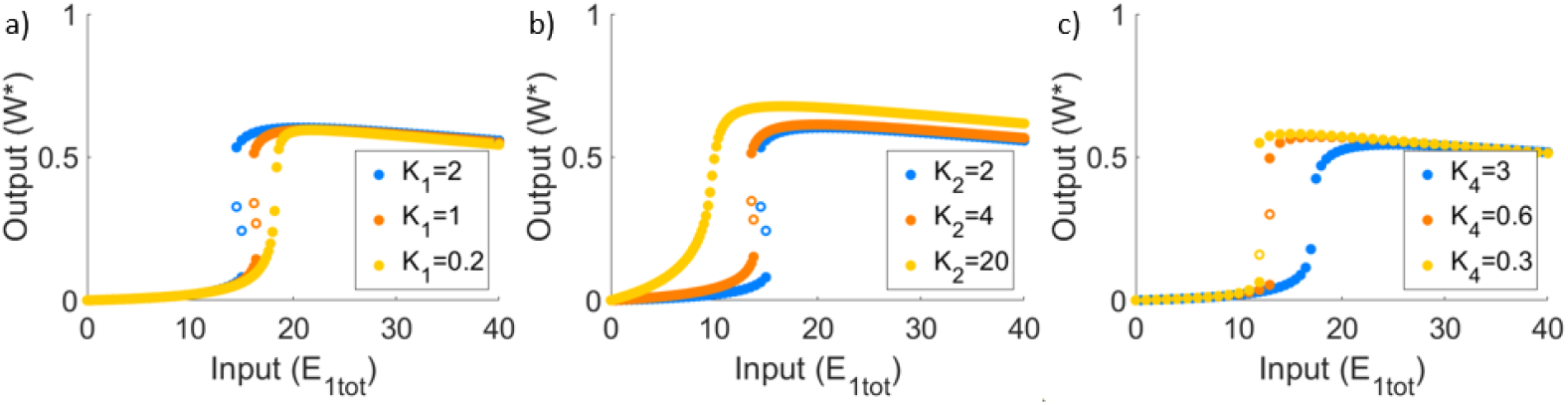
Michaelis constants can predispose PAR-i to bistability. Altering the Michaelis constants for a given value of *k*_4_ can take the system from monostability to bistability. Solid circles indicate stable solutions, while empty circles indicate unstable solutions. Parameters: all plots use *W*_*tot*_ = 100, *E*_2*tot*_ = 20, *a*_1_ = *d*_1_ = *k*_1_ = *a*_2_ = *d*_2_ = *k*_2_ = *a*_4_ = *d*_4_ = 1, *a*_3_ = 0.1, and *d*_3_ = 10. (a) *k*_4_ = 5, *a*_1_ = 1 (blue), *a*_1_ = 2 (orange), and *a*_1_ = 10 (yellow); (b) *k*_4_ = 5, *a*_2_ = 1 (blue), *a*_2_ = 0.5 (orange), and *a*_2_ = 0.1 (yellow); and (c) *k*_4_ = 2, *a*_4_ = 1 (blue), *a*_4_ = 5 (orange), *a*_4_ = 10. (yellow).

In common with the behaviour of PAR-d, we found that the catalytic constant, *k*_*i*_, is the only component rate constant in each Michaelis constant, *K*_*i*_, that exerts a qualitative influence on the shape of the system’s dose-response profile. In Fig 13b, for instance, we see that decreasing *k*_2_ increases the sensitivity of the profile, and may thereby predispose the system to bistability. On the other hand, increasing *k*_1_ increases sensitivity while also promoting monostability (see Fig 13a). In fact, we consistently found throughout our simulations that there is a very strong relationship between the values of *k*_1_ and *k*_4_, and the shape of the dose-response curve. In particular, for *k*_1_ = *k*_4_, this system is unable to achieve bistability, regardless of how the other parameters are altered.

**Figure 13.**
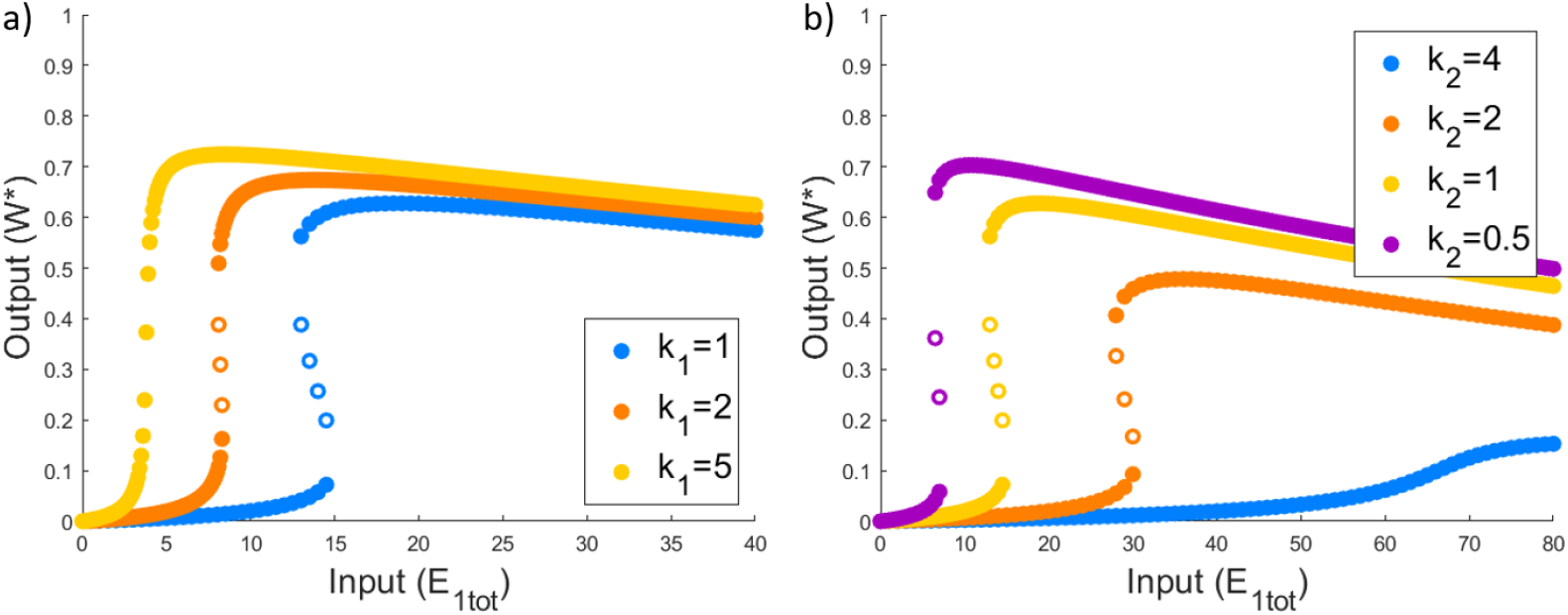
Low catalytic constants predispose PAR-i to bistability. Reducing the catalytic constants *k*_1_ and *k*_2_, for a given value of *k*_4_, can take the system from monostability to bistability. Solid circles indicate stable solutions and empty circles indicate unstable solutions. Parameters: *W*_*tot*_ = 100, *E*_2*tot*_ = 20, *a*_1_ = *d*_1_ = *k*_1_ = *a*_2_ = *d*_2_ = *k*_2_ = *a*_4_ = *d*_4_ = 1, *a*_3_ = 0.1, *d*_3_ = 10, and *k*_4_ = 10. (a) *a*_1_ = *d*_1_ = *k*_1_ = 1 (blue), *a*_1_ = *d*_1_ = *k*_1_ = 2 (orange), *a*_1_ = *d*_1_ = *k*_1_ = 5 (yellow); (b) *a*_2_ = *d*_2_ = *k*_2_ = 4 (blue), *a*_2_ = *d*_2_ = *k*_2_ = 2 (orange), *a*_2_ = *d*_2_ = *k*_2_ = 1 (yellow), *a*_2_ = *d*_2_ = *k*_2_ = 0.5 (purple).

As we show in Fig 14, increasing the total protein abundance, *W*_*tot*_ is also able to predispose the system to bistability.

**Figure 14.**
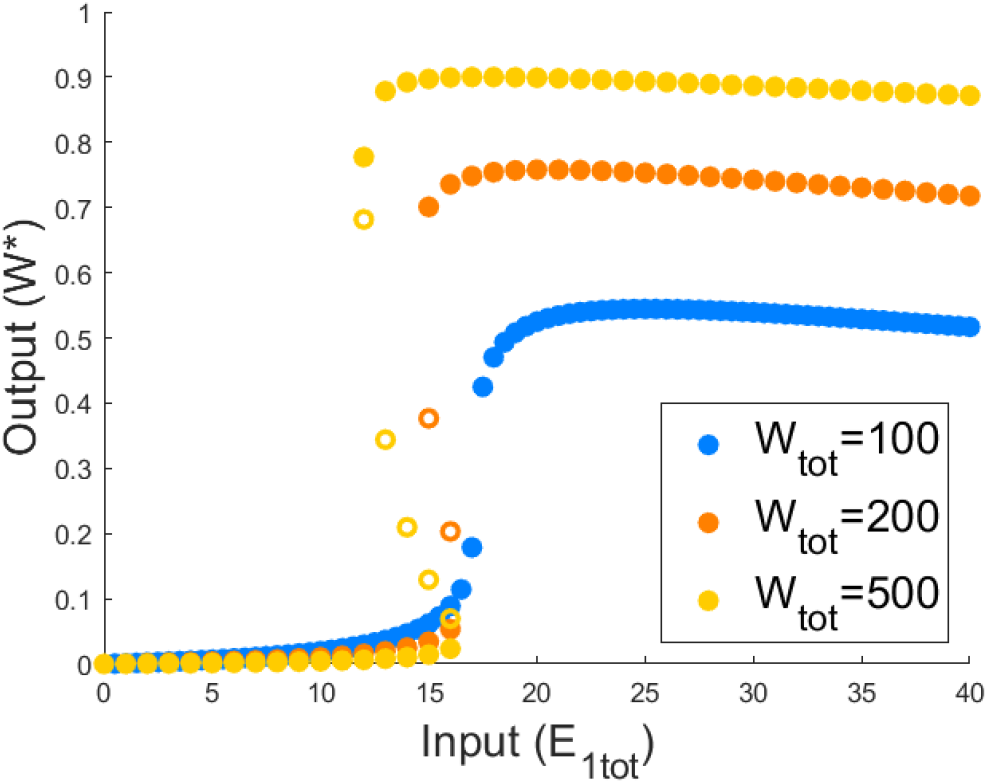
High total protein abundance predisposes PAR-i to bistability. Increasing the total abundance of protein substrate, *W*_*tot*_, for a given value of *k*_4_, can take the system from monostability to bistability. Solid circles indicate stable solutions and empty circles indicate unstable solutions. Parameters: *E*_2*tot*_ = 20, *a*_1_ = *d*_1_ = *k*_1_ = *a*_2_ = *d*_2_ = *k*_2_ = *a*_4_ = *d*_4_ = 1, *a*_3_ = 0.1, *d*_3_ = 10, and *k*_4_ = 2. We then set *W*_*tot*_ = 100 (blue), *W*_*tot*_ = 200 (orange), and *W*_*tot*_ = 500 (yellow).

Unlike the PAR-d model, but in common with the Michaelian model, PAR-i can only exhibit two-way (reversible) bistable switches, not one-way switches. As we illustrate in Fig 11, continued increases in the key parameter *k*_4_ do not culminate a one-way switch. This is to be expected, of course, since when *E*_1*tot*_ = 0 for this particular PAR mechanism, there is no possibility of a ‘forward’ reaction in which *W*\* is created, and thus no opportunity for a non-zero steady-state in *W*\*.

A further notable distinction between PAR-i and both the Michaelian and PAR-d models is the eventual decrease in system output with continued increases in input (*E*_1*tot*_). By analysing the abundances of intermediate complexes in this scenario, we find that after the maximum conversion from *W* to *W*\* is achieved (peak in *W*\* profile), and *E*_1*tot*_ continues to be increased, the third complex *C*_3_ continues to be produced without being consumed in the fourth reaction (due to insufficient *W*). This phenomenon is unique to the PAR-i model, and can be controlled by altering the value of 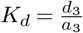. Indeed, we find that by increasing *K*_*d*_, we can mitigate this loss in maximum output (see Fig 15). Of course, inhibiting the formation of this complex in this manner also reduces the PAR contribution to the covalent modification cycle.

**Figure 15.**
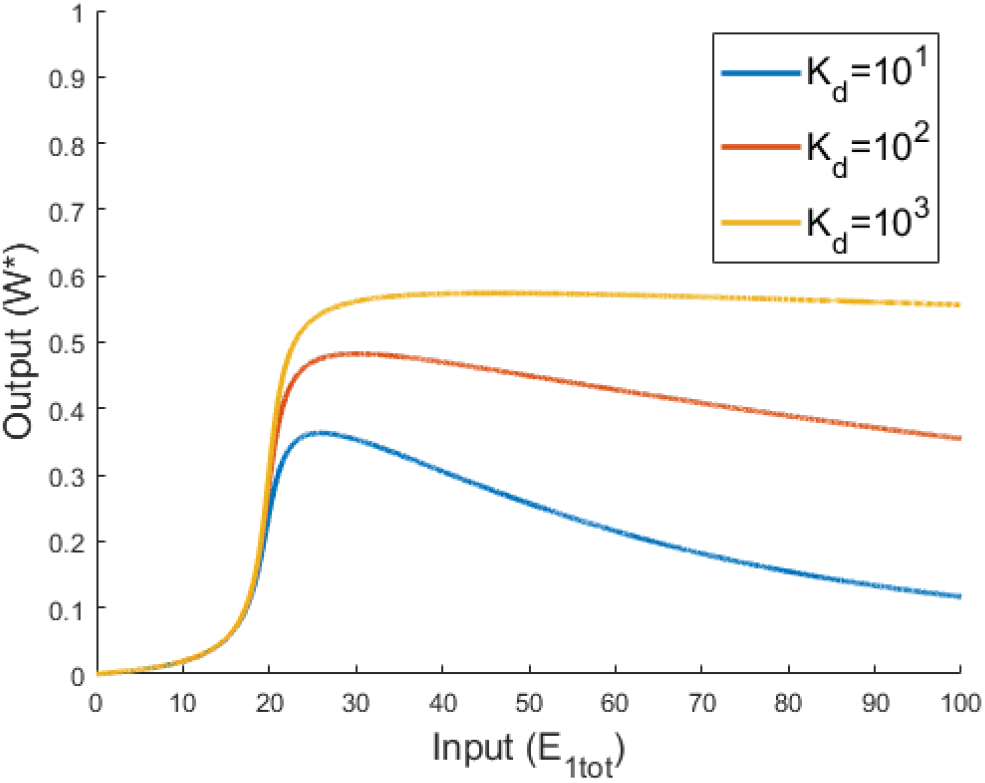
Effect of altering dissociation constant in PAR-i. We investigated increasing the value of *K*_*d*_ by decreasing the value of *a*_3_, although the same effect can be created by increasing *d*_3_. We found that by increasing the value of *K*_*d*_, the maximum abundance of modified substrate increases, and the removal of *W*\* after this maximum decreases. The parameters used are *W*_*tot*_ = 100. *E*_2*tot*_ = 20, *a*_1_ = *d*_1_ = *k*_1_ = *a*_2_ = *d*_2_ = *k*_2_ = *a*_4_ = *d*_4_ = *k*_4_ = 1, *d*_3_ = 10 with *a*_3_ = 1 (blue), *a*_3_ = 0.1 (red), and *a*_3_ = 0.01 (yellow).

### (c) Unlimited ultrasensitivity is difficult to achieve in complex-complete PAR models

Although ultrasensitive profiles could be obtained for both the PAR-d and the PAR-i models, it is important to consider whether or not the models support *unlimited* ultrasensitivity. In other words, can the slope of the highly sensitive (near-vertical) region of the dose-response profile be made arbitrarily steep in some parameter limit? Unlimited ultrasensitivity is of central importance to the implementation of robust perfect adaptation (RPA), and was incorporated into the influential RPA study by Ma et al. [13] using the Michaelian model discussed in the present work. Since the Michaelian model is capable of achieving unlimited ultrasensitivity in the parameter limit *K*_1_ → 0 and *K*_2_ → ∞ (*K*_1_ ≪ *W*_*tot*_ − *W*\* and *K*_2_ ≫ *W*\*), we examined the performance of our PAR-d and PAR-i models in this specific parameter regime. As we illustrate in Fig 16 we are unable to achieve unlimited ultrasensitivity using these conditions, for any choice of the remaining parameters in the model. In fact, for both the PAR-d and PAR-i models, as the limits *K*_1_ → 0 and *K*_2_ → ∞ are approached, the system exhibits a steep linear increase in *W*\* with *E*_1*tot*_ close to the origin.

**Figure 16.**
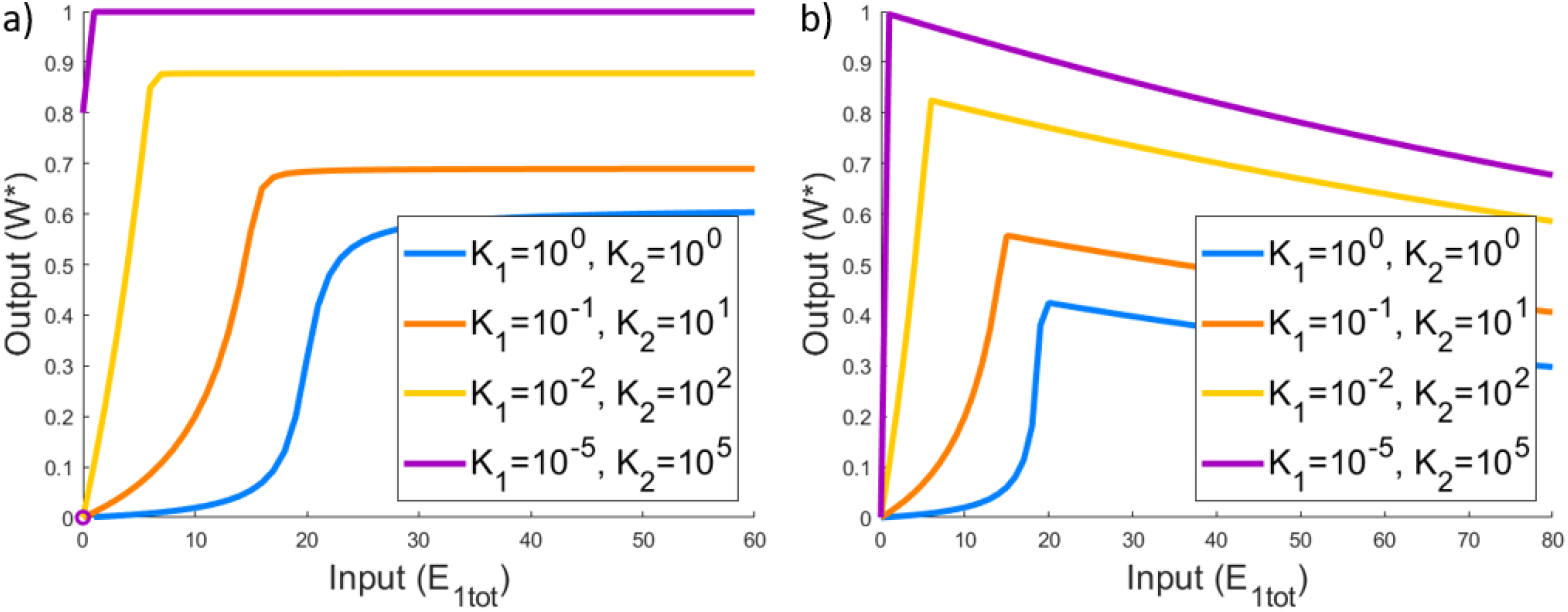
Parameter conditions corresponding to unlimited ultrasensitivity in the Michaelian model, applied to PAR-d and PAR-i. In the Michaelian model, unlimited ultrasensitivity obtains in the limit as *K*_1_ → 0 and *K*_2_ → ∞ (ie. *K*_1_ *W*_*tot*_ − *W*\* and *K*_2_ *W*\*). Here we consider the behaviour of the (a) PAR-d and (b) PAR-i models under these parametric conditions. Parameters: *W*_*tot*_ = 100, *E*_2*tot*_ = 20, *d*_1_ = *k*_1_ = *d*_2_ = *k*_2_ = 1; in addition: *a*_1_ = 1, *a*_2_ = 1 (blue); *a*_1_ = 10^−1^, *a*_2_ = 10^1^ (orange); *a*_1_ = 10^−2^, *a*_2_ = 10^2^ (yellow); *a*_1_ = 10^−5^, *a*_2_ = 10^5^ (purple). (a) *a*_3_ = 10^5^, *d*_3_ = *k*_3_ = 1, *K*_3_ = 10^−5^; (b) *a*_3_ = 0.1, *d*_3_ = 10, *a*_4_ = 10^5^, *d*_4_ = *k*_4_ = 1, *K*_*d*_ = 100, *K*_4_ = 10^−5^.

Our extensive numerical simulations revealed that the *only* parametric condition that yields unlimited ultrasensitivity in the PAR-d model is *a*_3_ → 0 (*K*_3_ → ∞), along with either *K*_1_ → 0 and *K*_2_ → 0 (Fig 17a) or *W*_*tot*_ ≫ *E*_2*tot*_, *E*_1*tot*_ (Fig 17b). We note that the first of these conditions (*a*_3_ → 0) corresponds to the limit in which the PAR-d mechanism approaches the Goldbeter-Koshland model [46] - ie. a simple covalent-modification cycle *without* positive autoregulation (PAR). In addition, the second set of conditions (*K*_1_ → 0 and *K*_2_ → 0 or *W*_*tot*_ ≫ *E*_2*tot*_, *E*_1*tot*_) corresponds to parameter regime in which Goldbeter and Koshland [46] achieved unlimited ultrasensitivity via the zero-order mechanism. In other words PAR-d can only exhibit unlimited ultrasensitivity once the PAR contribution is removed. PAR-d, in its essence, appears to be incapable of unlimited ultrasensitivity.

**Figure 17.**
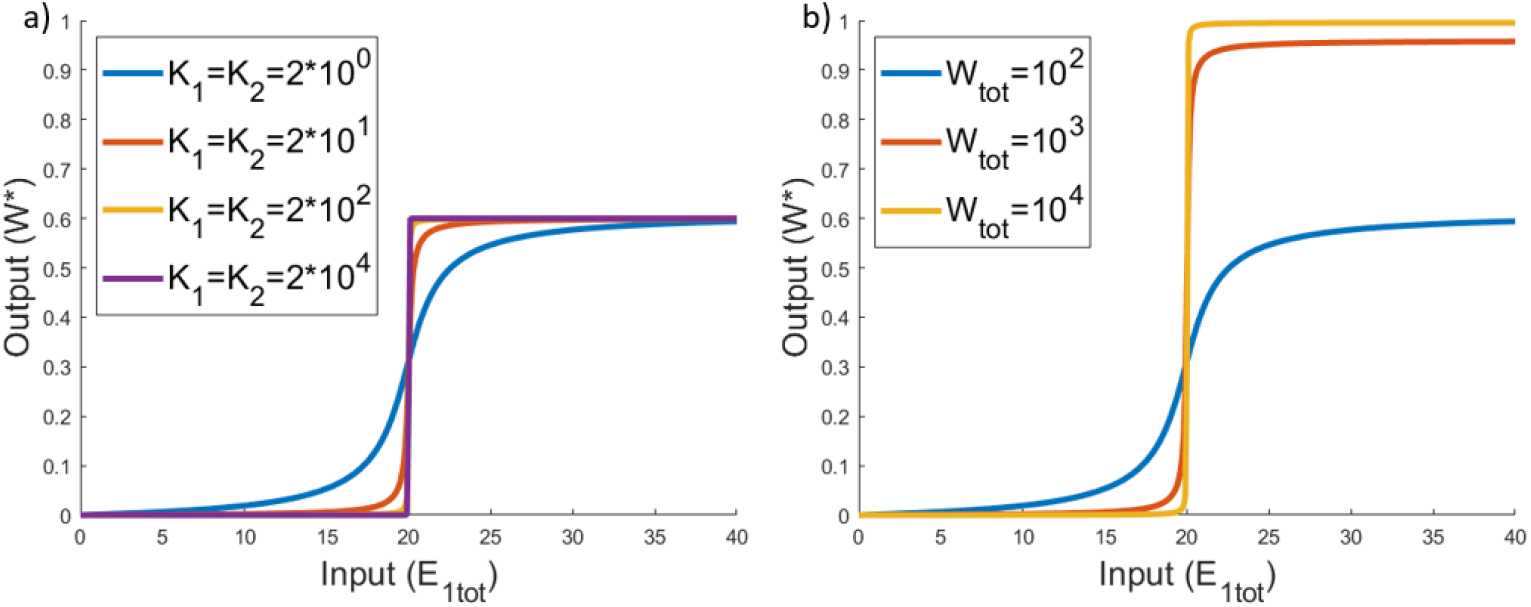
Unlimited Ultrasensitivity in PAR-d. Unlimited ultrasensitivity can only obtain in PAR-d in the limit as *a*_3_ → 0, whereby the positive autoregulation is removed from the model, and *K*_1_, *K*_2_ ≪ *W*_*tot*_, which corresponds to the conditions required for zero-order ultrasensitivity. These conditions can be realized by (a) decreasing *K*_1_ and *K*_2_, or (b) increasing *W*_*tot*_. Parameters: *W*_*tot*_ = 100, *E*_2*tot*_ = 20, *a*_1_ = *d*_1_ = *k*_1_ = *a*_2_ = *d*_2_ = *k*_2_ = *d*_3_ = *k*_3_ = 1, and *a*_3_ = 10^−10^. (a) *a*_1_ = *a*_2_ = 1 (blue), *a*_1_ = *a*_2_ = 10^1^ (red), *a*_1_ = *a*_2_ = 10^2^ (yellow), and *a*_1_ = *a*_2_ = 10^4^ (purple); (b) *W*_*tot*_ = 10^2^ (blue), *W*_*tot*_ = 10^3^ (red), and *W*_*tot*_ = 10^4^ (yellow). Almost identical profiles can be achieved in PAR-i by setting *a*_3_ = 10^−10^, *d*_3_ = 10, and *a*_4_ = *d*_4_ = *k*_4_ = 1 to remove the effect of PAR (not shown).

Unlimited ultrasensivity was likewise very difficult to find in PAR-i, although it is possible under very specific conditions. Similar to the situation with PAR-d, unlimited ultrasensitivity may be realised in the parameter limit by which the Goldbeter-Koshland model [46] is recovered from the PAR-i model. This gives rise to dose-response profiles which are identical to Fig 17.

However PAR-i is also able to create unlimited ultrasensitivity in one additional set of conditions, where PAR remains present in the model. For this, we consistently found that we require *k*_1_ ≈ *k*_2_ ≈ *k*_4_. When this condition holds, unlimited ultrasensitivity may then be achieved by taking the limit as *K*_1_ → 0 and *K*_2_ → 0 (Fig 18a), and/or in the limit as *W*_*tot*_ → 0 (Fig 18b).

**Figure 18.**
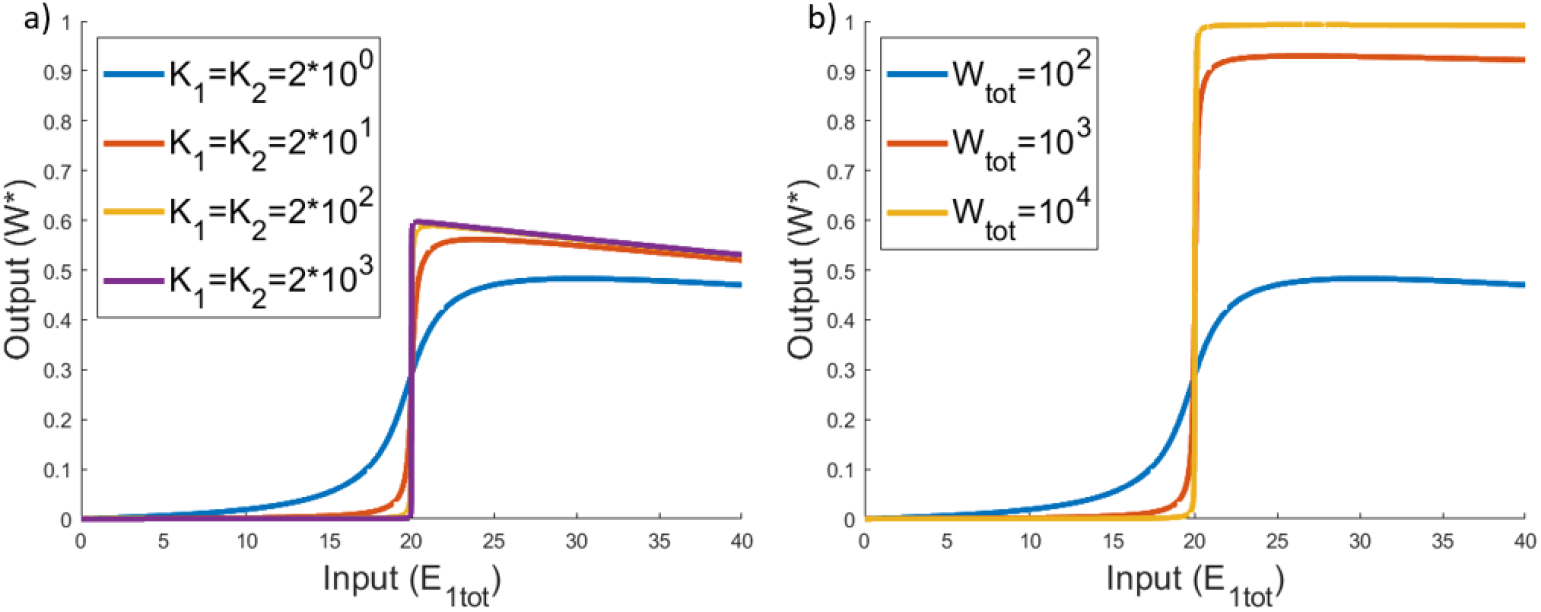
Unlimited Ultrasensitivity in PAR-i. In this figure we create conditions on PAR-i under which we can observe unlimited ultrasensitivity without sacrificing the effect of the PAR mechanism. To achieve these plots, we apply the same conditions required for unlimited ultrasensitivity in a reversible covalent-modification cycle as Fig 17. However it is still evident that the PAR mechanism has some effect on the profile, as demonstrated by the distinct decline in modified substrate after the maximum conversion is achieved. A behaviour which is unique to this PAR mechanism. These profiles are achieved by setting *W*_*tot*_ = 100, *E*_2*tot*_ = 20, *a*_1_ = *d*_1_ = *k*_1_ = *a*_2_ = *d*_2_ = *k*_2_ = *a*_4_ = *d*_4_ = *k*_4_ = 1, *a*_3_ = 0.1, and *d*_3_ = 10. We then set a) *a*_1_ = *a*_2_ = 1 (blue), *a*_1_ = *a*_2_ = 10^1^ (red), *a*_1_ = *a*_2_ = 10^2^ (yellow), and *a*_1_ = *a*_2_ = 10^3^ (purple) and b) *W*_*tot*_ = 10^2^ (blue), *W*_*tot*_ = 10^3^ (red), and *W*_*tot*_ = 10^4^ (yellow)

## 3. Conclusions

The Michaelis-Menten equation, and variations thereon, has appeared almost ubiquitously in mathematical descriptions of enzyme-mediated chemical reactions for more than 100 years [78,79] - often quite indiscriminately, with little or no consideration of whether it is able to truly recapitulate the behaviour of the (potentially much more complicated) system being modelled. In particular, it has been used in highly-influential work [13] to suggest that positive autoregulation (PAR) of a covalent-modification cycle can generate unlimited ultrasensitivity, and as a consequence, robust perfect adaptation (RPA) when suitably embedded in certain network topologies. Here we ask: Can a Michaelian description of such a system accurately represent the detailed molecular mechanisms that must be present in a ‘real’ collection of interacting molecules, which could, in principle, be quite intricate and complicated? And can it truly recapitulate the range of qualitative behaviours afforded by more detailed mass-action descriptions that account for all molecular interactions? Our answer, based on two detailed mass-action models which carefully account for the specific molecular mechanisms through which PAR is actually transacted, and explicitly account for all intermediate molecular species - a framework we call ‘*complex complete*’ - is a resounding *no*.

Although our two complex-complete PAR models (PAR-d and PAR-i) are relatively simple, they exhibit functionally important differences from their Michaelian counterpart, and clearly illustrate that *mechanism matters*. Indeed, although the Michaelian model of PAR has been studied for its capacity to exhibit unlimited ultrasensitivity for the purposes of RPA, we found that our more realistic models that account for the specific mechanism involved were not particularly prone to unlimited ultrasensitivity at all, and required very carefully-orchestrated parametric conditions in order to be able to increase the sensitivities of their dose-response profiles indefinitely. For one of the two mechanisms (PAR-d), unlimited ultrasensitivity could *only* be obtained in the parameter limit corresponding to the complete removal of positive autoregulation from the covalent modification cycle. In any event, neither of our complex-complete models could achieve ultrasensitivity in the parametric circumstances suggested by the Michaelian model (very small Michaelis constant for the ‘forward’ reaction, and very large Michaelis constant for the ‘reverse’ reaction).

Both complex-complete PAR models were prone to bistability, however - a qualitative response that is also realised by the Michaelian description. But here again, behaviour of the complex-complete models departs from that of the Michaelian model in significant ways. The PAR-d model - by far the more prone to bistability of the two new models - may readily be shown to exhibit both two-way (reversible) bistable switches *and* one-way (irreversible) switches. Moreover, a full range of qualitative responses - from a monostable ultrasensitive profile, to two-way switches, through to one-way switches - could be generated in the PAR-d model *for a single set of Michaelis constants*, the specific response being determined in each case by *other* component parameters in the model. In contrast, the Michaelis constants alone specify the qualitative nature of the dose-response curve for the Michaelian model. The PAR-i model was shown to be much less prone to bistability than the PAR-d model, and could only exhibit two-way (reversible) switches - a response that is entirely driven by the catalytic rate constant associated with the autoregulatory enzymatic reaction.

Our study suggests that more caution should be exercised in the use of the Michaelis-Menten equation to model enzyme-mediated reactions, particularly in the context of complex signalling networks, and particularly when additional regulations (such as autoregulation) are present. While it is clear that Michaelian models vastly simplify the mathematical analysis of enzyme-mediated reaction networks, where complex-complete models can quickly become very large and complicated, these simpler models can only make useful predictions if they can, in fact, recapitulate the basic qualitative features of their more detailed counterparts.

We note in closing that complex-complete mass-action models, as we propose here, constitute polynomial dynamical systems whose steady-state solutions form algebraic varieties. Significant progress has been made in recent years in the application of Gröbner basis methods to the study of these kinds of models (see for example [80]). Nevertheless, since Gröbner bases can quickly become large and expensive to compute as the underlying models become more complicated, numerical simulations may still be required in many cases to support the analysis of mechanism-based models of complex networks.

## 4. Methods

Numerical simulations and root finding methods were all undertaken using standard packages available in Matlab (ODE45, ODE23s, roots, and fsolve). Total number of possible steady-states in the interval [0,*W*_*tot*_] were checked by Gröbner basis calculations (and subsequent numerical examination in Matlab) available in the open-source package Singular (https://www.singular.uni-kl.de/). All code available from the authors upon reasonable request.

## Data Accessibility

This article has no additional data.

## Authors’ Contributions

CJS: conceptualization, formal analysis, methodology, software, visualisation, writing-original draft preparation. RPA: conceptualization, methodology, writing – review and editing, supervision.

## Acknowledgements

Robyn P. Araujo is the recipient of an Australian Research Council (ARC) Future Fellowship (project number FT190100645) funded by the Australian Government.

